## Supplementary material for "Coronavirus testing indicates transmission risk increases along wildlife supply chains for human consumption in Viet Nam, 2013-2014": S1 R code: S1_rcode.html

#### 5/29/2020

### Load Data

Load data from dryad repository:

```
temp <- tempfile()
download.file("https://datadryad.org/stash/share/pk3wVUxFNzTuCYZ9t8haKRPmx7V8YhTDBuHpG8JJ9kU",temp, mode="wb")
#download.file("https://doi.org/10.5061/dryad.7h44j0zrj",temp, mode="wb") 
unzip(temp, "VietnamWithInterpretations_20200323_dryad.csv")
H1 <- read.csv("VietnamWithInterpretations_20200323_dryad.csv")
head(H1)
```

```
##   Country                             SiteName StateProv
## 1 Vietnam Bat roost and quano farm - Soc Trang Soc Trang
## 2 Vietnam Bat roost and quano farm - Soc Trang Soc Trang
## 3 Vietnam Bat roost and quano farm - Soc Trang Soc Trang
## 4 Vietnam Bat roost and quano farm - Soc Trang Soc Trang
## 5 Vietnam Bat roost and quano farm - Soc Trang Soc Trang
## 6 Vietnam Bat roost and quano farm - Soc Trang Soc Trang
##                                         EventName
## 1 Bat roost and quano farm - Soc Trang-2013-01-10
## 2 Bat roost and quano farm - Soc Trang-2013-01-10
## 3 Bat roost and quano farm - Soc Trang-2013-01-10
## 4 Bat roost and quano farm - Soc Trang-2013-01-10
## 5 Bat roost and quano farm - Soc Trang-2013-01-10
## 6 Bat roost and quano farm - Soc Trang-2013-01-10
##                     DomesticAnimals      AnimalID TaxaGroup
## 1 pigs, ducks, cats, chickens, dogs VN13M0199 F2R      Bats
## 2 pigs, ducks, cats, chickens, dogs VN13M0199 F2R      Bats
## 3 pigs, ducks, cats, chickens, dogs VN13M0200 F1R      Bats
## 4 pigs, ducks, cats, chickens, dogs VN13M0200 F1R      Bats
## 5 pigs, ducks, cats, chickens, dogs VN13M0200 F2R      Bats
## 6 pigs, ducks, cats, chickens, dogs VN13M0200 F2R      Bats
##   SpeciesScientificName       CommonName
## 1            Chiroptera Unidentified Bat
## 2            Chiroptera Unidentified Bat
## 3            Chiroptera Unidentified Bat
## 4            Chiroptera Unidentified Bat
## 5            Chiroptera Unidentified Bat
## 6            Chiroptera Unidentified Bat
##                      CommonNameFieldMorphology        IDCertainty      Order
## 1 Unidentified Bat Within The Chiroptera Order field ID uncertain CHIROPTERA
## 2 Unidentified Bat Within The Chiroptera Order field ID uncertain CHIROPTERA
## 3 Unidentified Bat Within The Chiroptera Order field ID uncertain CHIROPTERA
## 4 Unidentified Bat Within The Chiroptera Order field ID uncertain CHIROPTERA
## 5 Unidentified Bat Within The Chiroptera Order field ID uncertain CHIROPTERA
## 6 Unidentified Bat Within The Chiroptera Order field ID uncertain CHIROPTERA
##   Family Genus SampleDate    SpecimenID SpecimenType         TestType
## 1   NULL  NULL    1/14/13 VN13M0199 F2R        feces Conventional PCR
## 2   NULL  NULL    1/14/13 VN13M0199 F2R        feces Conventional PCR
## 3   NULL  NULL    1/14/13 VN13M0200 F1R        feces Conventional PCR
## 4   NULL  NULL    1/14/13 VN13M0200 F1R        feces Conventional PCR
## 5   NULL  NULL    1/14/13 VN13M0200 F2R        feces Conventional PCR
## 6   NULL  NULL    1/14/13 VN13M0200 F2R        feces Conventional PCR
##   TestRequested              TestRequestedProtocol ConfirmationResult Sequence
## 1 Coronaviruses Modified Watanabe et al, RdRp gene           Negative     NULL
## 2 Coronaviruses              Quan et al, RdRp gene           Negative     NULL
## 3 Coronaviruses              Quan et al, RdRp gene           Negative     NULL
## 4 Coronaviruses Modified Watanabe et al, RdRp gene           Negative     NULL
## 5 Coronaviruses Modified Watanabe et al, RdRp gene           Negative     NULL
## 6 Coronaviruses              Quan et al, RdRp gene           Negative     NULL
##   GenbankAccessionNumber Virus VirusGroup
## 1                   NULL  NULL       NULL
## 2                   NULL  NULL       NULL
## 3                   NULL  NULL       NULL
## 4                   NULL  NULL       NULL
## 5                   NULL  NULL       NULL
## 6                   NULL  NULL       NULL
```

```
colnames(H1)
```

```
##  [1] "Country"                   "SiteName"                 
##  [3] "StateProv"                 "EventName"                
##  [5] "DomesticAnimals"           "AnimalID"                 
##  [7] "TaxaGroup"                 "SpeciesScientificName"    
##  [9] "CommonName"                "CommonNameFieldMorphology"
## [11] "IDCertainty"               "Order"                    
## [13] "Family"                    "Genus"                    
## [15] "SampleDate"                "SpecimenID"               
## [17] "SpecimenType"              "TestType"                 
## [19] "TestRequested"             "TestRequestedProtocol"    
## [21] "ConfirmationResult"        "Sequence"                 
## [23] "GenbankAccessionNumber"    "Virus"                    
## [25] "VirusGroup"
```

### Clean up data/variables

#### Variable names

```
names(H1) <- to_snake_case(names(H1))
```

#### Dates

```
H1$sample_date<-as.character(H1$sample_date)
H1$sample_date<-as.POSIXct(strptime(H1$sample_date, "%m/%d/%y", tz="Asia/bangkok"))
H1$sample_month<-month(H1$sample_date)
```

Note: The sample dates that are earlier than 2011 correspond to archived samples that were obtained from Tam Dao Bear rescue center

#### Creating primary interface group

Establish primary interface groups based on site names.

```
H1$primary_interface_group<-NA
H1$primary_interface_group<-as.character(rep("captive facilities", n=nrow(H1)))
H1$primary_interface_group[is.element(H1$site_name, c("Rat market CT - Dong Thap","Rat market LVO - Dong Thap", "Rat market LVU - Dong Thap", "Rat market MX1 - Soc Trang", "Rat market MX2 - Soc Trang", "Rat market N5(1) - Soc Trang", "Rat market N5(2) - Soc Trang", "Rat market SD - Dong Thap"))] <-'trader' 
H1$primary_interface_group[is.element(H1$site_name, c("Rat market CLC1 - Dong Thap", "Rat market CLC2 - Dong Thap", "Rat market CLC3 - Dong Thap", "Rat market CLD - Dong Thap", "Rat market HND - Dong Thap", "Rat market HNT - Dong Thap", "Rat market TB1 - Dong Thap", "Rat market TB2 - Dong Thap", "Rat market TH - Dong Thap", "Rat market TM1 - Dong Thap", "Rat market TM2 - Dong Thap", "Rat market TN - Dong Thap", "Rat market TT - Soc Trang", "Wet market (Whole sale sellers) - Soc Trang"))] <-'large market' 
H1$primary_interface_group[is.element(H1$site_name, c("CPCP civet restaurant confiscation","Rat restaurant ST - Soc Trang", "Restaurant - Soc Trang", "Lam Dong FPD"))] <-'restaurant' 
H1$primary_interface_group[is.element(H1$site_name, c("Bat pagoda - Soc Trang"))] <-'natural roost' 
H1$primary_interface_group[is.element(H1$site_name, c("Bat roost and quano farm - Soc Trang", "Bat roost and quano farm CL - Dong Thap", "Bat roost and quano farm CLD2 - Soc Trang", "Bat roost and quano farm CLD3 - Soc Trang", "Bat roost and quano farm CLD5 - Soc Trang", "Bat roost and quano farm CLD6 - Soc Trang", "Bat roost and quano farm CLD7 - Soc Trang", "Bat roost and quano farm CT - Dong Thap", "Bat roost and quano farm HN - Dong Thap", "Bat roost and quano farm LP - Soc Trang", "Bat roost and quano farm LV - Dong Thap", "Bat roost and quano farm TM - Dong Thap", "Bat roost and quano farm CLD1 - Soc Trang", "Bat roost and quano farm CLD4 - Soc Trang", "Bat roost and quano farm MX - Soc Trang", "Bat roost and quano farm N5(1) - Soc Trang", "Bat roost and quano farm N5(2) - Soc Trang"))] <-'bat guano farm' 
H1$primary_interface_group[is.element(H1$site_name, c("Farm 1 -  Bien Hoa - DN", "Farm 10 -  Vinh Cuu - DN", "Farm 12 - Nhon Trach - DN", "Farm 13 - Nhon Trach - DN", "Farm 14 - Long Thanh - DN", "Farm 15 - Long Thanh - DN", "Farm 16 - Long Thanh - DN", "Farm 17 - Long Thanh - DN", "Farm 18 - Cam My - DN", "Farm 19 - Cam My - DN", "Farm 2 -  Bien Hoa - DN", "Farm 20 - Cam My - DN", "Farm 21 - Xuan Loc - DN", "Farm 22 - Xuan Loc - DN", "Farm 23 - Xuan Loc - DN", "Farm 24 - Xuan Loc - DN", "Farm 25 - Long Khanh - DN", "Farm 26 - Xuan Loc - DN", "Farm 28 - Dinh Quan - DN", "Farm 29 - Tan Phu - DN", "Farm 3 -  Bien Hoa - DN", "Farm 4 -  Thong Nhat - DN", "Farm 5 -  Thong Nhat - DN", "Farm 6 -  Trang Bom - DN", "Farm 8 -  Vinh Cuu - DN", "Farm 9 -  Vinh Cuu - DN", "Wildlife farm 1 - Dong Nai", "Wildlife farm 3 - Dong Nai", "Wildlife farm 4 - Dong Nai"))] <-'wildlife farm' 
H1$primary_interface_group <- as.factor(H1$primary_interface_group)
H1$primary_interface_group <- factor(H1$primary_interface_group, levels = c("trader", "large market", "restaurant", "natural roost","bat guano farm","wildlife farm","captive facilities"))
```

#### Cleaning genus

Add family for some missing genus in the data (couple squirrels and unidentified viverridae). Also added the captive facility for the bears and macaques, even though they tested negative (avoid missing values)

```
H1$new_genus<-NA
H1$new_genus<-as.character(H1$genus)
H1<-H1 %>% 
  mutate(new_genus=replace(new_genus,H1$genus=="NULL" & species_scientific_name=="Viverridae","Viverridae"))
H1<-H1 %>% 
  mutate(new_genus=replace(new_genus,H1$genus=="NULL" & species_scientific_name=="Sciuridae","Sciuridae"))
H1<-H1 %>% 
  mutate(new_genus=replace(new_genus,H1$genus=="NULL" & species_scientific_name=="Chiroptera","Microchiroptera"))
H1<-H1 %>% 
  mutate(new_genus=replace(new_genus,primary_interface_group=="trader" & family=="MURIDAE","Field Rats"))
H1<-H1 %>% 
  mutate(new_genus=replace(new_genus,primary_interface_group=="large market" & family=="MURIDAE","Field Rats"))
H1<-H1 %>% 
  mutate(new_genus=replace(new_genus,primary_interface_group=="restaurant" & family=="MURIDAE","Field Rats"))
```

#### Create variable that lists viruses associated with each animal id to determine coinfection status

Each row in the data set is a test and virus sequence result so need to add a field that summarizes all viruses found in a single individual to establish co-infection status

```
tempH1<-as.data.frame(H1)
tempH1$virus<-as.character(tempH1$virus)
tempH1$virus[tempH1$virus == "NULL"] <- NA #replace NULL with NA

temp1 <- tempH1 %>%
  drop_na(virus) %>% 
  group_by(animal_id) %>%
  summarize(virus_ind = (paste0(unique(virus_group), collapse=" | "))) %>% 
  as.data.frame()

temp2 <- tempH1 %>%
  group_by(animal_id, specimen_id) %>%
  summarize(virus_specimen = paste(unique(virus_group), collapse=" | "))%>%   as.data.frame()

temp3<-full_join(temp1, temp2, by="animal_id")

temp4<-subset(temp3, select=-animal_id)

H1<-full_join(temp4, H1, by="specimen_id")
```

#### Define wet and dry seasons based on sampling date

The annual wet season in southern Vietnam occurs from May 1st through November 30th, and the dry season from December 1st through April 30th.

```
H1$hseason <- NA
H1$hseason <- ifelse(H1$sample_month %in% c(12,1:4), "Dry","Wet")
table(H1$sample_month,H1$hseason)
```

```
##     
##       Dry  Wet
##   1   621    0
##   3   788    0
##   4    70    0
##   10    0 2793
##   11    0   74
```

#### Select just rodents and bats for futher analysis

```
H1<-subset(H1, is.element(taxa_group,c("Bats","Rodents & Shrews")))
H1<-droplevels(H1)
```

### Materials and methods

#### Sampling locations

Sampling site description by interface type and province:

```
H1 %>% 
  group_by(primary_interface_group) %>%
  summarize(site_counts=length(unique(site_name)))
```

```
## # A tibble: 6 x 2
##   primary_interface_group site_counts
##   <fct>                         <int>
## 1 trader                            8
## 2 large market                     14
## 3 restaurant                        2
## 4 natural roost                     1
## 5 bat guano farm                   17
## 6 wildlife farm                    28
```

```
H1 %>% 
  group_by(primary_interface_group, state_prov) %>%
  summarize(site_counts=length(unique(site_name)))
```

```
## # A tibble: 9 x 3
## # Groups:   primary_interface_group [6]
##   primary_interface_group state_prov site_counts
##   <fct>                   <fct>            <int>
## 1 trader                  Dong Thap            4
## 2 trader                  Soc Trang            4
## 3 large market            Dong Thap           12
## 4 large market            Soc Trang            2
## 5 restaurant              Soc Trang            2
## 6 natural roost           Soc Trang            1
## 7 bat guano farm          Dong Thap            5
## 8 bat guano farm          Soc Trang           12
## 9 wildlife farm           Dong Nai            28
```

Timing of sampling:

```
H1 %>% 
  group_by(hseason) %>%
  summarize(site_count=length(unique(site_name)))
```

```
## # A tibble: 2 x 2
##   hseason site_count
##   <chr>        <int>
## 1 Dry             30
## 2 Wet             41
```

```
min(H1$sample_date)
```

```
## [1] "2013-01-11 +07"
```

```
max(H1$sample_date)
```

```
## [1] "2014-03-29 +07"
```

Number of visits per site (and per season):

```
H1 %>% 
  group_by(site_name) %>%
  summarize(site_visits=length(unique(event_name)))
```

```
## # A tibble: 70 x 2
##    site_name                                 site_visits
##    <fct>                                           <int>
##  1 Bat pagoda - Soc Trang                              3
##  2 Bat roost and quano farm - Soc Trang                1
##  3 Bat roost and quano farm CL - Dong Thap             1
##  4 Bat roost and quano farm CLD1 - Soc Trang           1
##  5 Bat roost and quano farm CLD2 - Soc Trang           1
##  6 Bat roost and quano farm CLD3 - Soc Trang           1
##  7 Bat roost and quano farm CLD4 - Soc Trang           1
##  8 Bat roost and quano farm CLD5 - Soc Trang           1
##  9 Bat roost and quano farm CLD6 - Soc Trang           1
## 10 Bat roost and quano farm CLD7 - Soc Trang           1
## # … with 60 more rows
```

```
H1 %>% 
  group_by(site_name,hseason) %>% 
  summarize(vists=length(unique(sample_date)))
```

```
## # A tibble: 71 x 3
## # Groups:   site_name [70]
##    site_name                                 hseason vists
##    <fct>                                     <chr>   <int>
##  1 Bat pagoda - Soc Trang                    Dry         5
##  2 Bat pagoda - Soc Trang                    Wet         1
##  3 Bat roost and quano farm - Soc Trang      Dry         1
##  4 Bat roost and quano farm CL - Dong Thap   Wet         1
##  5 Bat roost and quano farm CLD1 - Soc Trang Wet         1
##  6 Bat roost and quano farm CLD2 - Soc Trang Wet         1
##  7 Bat roost and quano farm CLD3 - Soc Trang Wet         1
##  8 Bat roost and quano farm CLD4 - Soc Trang Wet         1
##  9 Bat roost and quano farm CLD5 - Soc Trang Wet         1
## 10 Bat roost and quano farm CLD6 - Soc Trang Wet         1
## # … with 61 more rows
```

In most cases, there was only one visit per site, except for the bat pagoda. This should call for caution when looking at seasonal effect, as site and season are confounded (so season effect may just be a result of the timing of sampling across the different sites). Only the Soc Trang bat pagoda had sampling in two seasons and is suitable for assessment of seasonal effect.

#### Animal Sampling

Summary tables clarifying what sampling protocol was used in each interface/site.

```
H1 %>% 
select(primary_interface_group,specimen_type) %>% 
  group_by(primary_interface_group) %>% 
  count(specimen_type) %>% 
  pivot_wider(names_from = specimen_type,values_from = n)
```

```
## # A tibble: 6 x 12
## # Groups:   primary_interface_group [6]
##   primary_interfa… brain  lung `oral swab` `small intestin… feces kidney
##   <fct>            <int> <int>       <int>            <int> <int>  <int>
## 1 trader              50    94         308              232    NA     NA
## 2 large market        90   102         589              443    26    122
## 3 restaurant          NA   170         239              191     4     NA
## 4 natural roost       NA    NA          30               NA    90     NA
## 5 bat guano farm      NA    NA          NA               NA   624     NA
## 6 wildlife farm       NA    NA          NA               NA   796     NA
## # … with 5 more variables: `rectal swab` <int>, `urine/urogenital swab` <int>,
## #   spleen <int>, urine <int>, `environmental sample` <int>
```

```
H1 %>% 
select(site_name,specimen_type) %>% 
  group_by(site_name) %>% 
  count(specimen_type) %>% 
  pivot_wider(names_from = specimen_type,values_from = n)
```

```
## # A tibble: 70 x 12
## # Groups:   site_name [70]
##    site_name feces `oral swab` `rectal swab` urine `environmental …
##    <fct>     <int>       <int>         <int> <int>            <int>
##  1 Bat pago…    90          30            36     4               NA
##  2 Bat roos…    16          NA            NA    NA               NA
##  3 Bat roos…    19          NA            NA    NA               NA
##  4 Bat roos…    12          NA            NA    NA               NA
##  5 Bat roos…    20          NA            NA    NA               NA
##  6 Bat roos…    14          NA            NA    NA               NA
##  7 Bat roos…    10          NA            NA    NA               NA
##  8 Bat roos…    24          NA            NA    NA               NA
##  9 Bat roos…     6          NA            NA    NA               NA
## 10 Bat roos…    10          NA            NA    NA               NA
## # … with 60 more rows, and 6 more variables: `urine/urogenital swab` <int>,
## #   kidney <int>, `small intestine` <int>, brain <int>, lung <int>,
## #   spleen <int>
```

### Results

#### Detection of coronavirus by animal taxa and interface

##### Paragraph 1 overview of specimens and animals

Number of samples:

```
length(unique(H1$specimen_id))
```

```
## [1] 2164
```

Number of animals:

```
length(unique(H1$animal_id)) #ID assigned in the field
```

```
## [1] 1506
```

Break-down of animal number and proportion of positive by taxonomic group:

```
H1 %>%
  group_by(taxa_group) %>%
  summarize(ind_count=length(unique(animal_id)),
            ind_count_pos=length(unique(animal_id[confirmation_result=="Positive"])),
            percent_pos = ind_count_pos/ind_count*100)
```

```
## # A tibble: 2 x 4
##   taxa_group       ind_count ind_count_pos percent_pos
##   <fct>                <int>         <int>       <dbl>
## 1 Bats                   375           238        63.5
## 2 Rodents & Shrews      1131           266        23.5
```

Distribution of samples and animals by province

```
H1 %>% 
  group_by(state_prov) %>%
  summarize(ind_count=length(unique(animal_id)), site_count=length(unique(site_name)))
```

```
## # A tibble: 3 x 3
##   state_prov ind_count site_count
##   <fct>          <int>      <int>
## 1 Dong Nai         429         28
## 2 Dong Thap        565         21
## 3 Soc Trang        512         21
```

##### S1 Table

Summary of all testing results by genus, interface, sub-interface, sample types, number specimens tested, number specimens tested positives, %positive and viral species.

```
S1Table <- H1 %>%
  group_by(new_genus, primary_interface_group, specimen_type) %>%
  summarize(site_count = length(unique(site_name)),
            ind_count_pos=length(unique(animal_id[confirmation_result=="Positive"])),
            ind_count=length(unique(animal_id)),
            percent_pos = ind_count_pos/ind_count*100,
            IndivLevel= paste0(round(percent_pos,digits = 1), "% (",ind_count_pos,"/",ind_count,")"),
            viruses = paste(unique(virus_specimen), collapse=", "))
S1Table
```

```
## # A tibble: 32 x 9
## # Groups:   new_genus, primary_interface_group [10]
##    new_genus primary_interfa… specimen_type site_count ind_count_pos ind_count
##    <chr>     <fct>            <fct>              <int>         <int>     <int>
##  1 Cynopter… natural roost    oral swab              1             0         1
##  2 Cynopter… natural roost    rectal swab            1             0         2
##  3 Field Ra… trader           brain                  1             1        25
##  4 Field Ra… trader           lung                   4             6        47
##  5 Field Ra… trader           oral swab              8            28       154
##  6 Field Ra… trader           small intest…          7            14       116
##  7 Field Ra… large market     brain                  5             4        45
##  8 Field Ra… large market     feces                  1             0        13
##  9 Field Ra… large market     kidney                 8             3        61
## 10 Field Ra… large market     lung                   3             4        51
## # … with 22 more rows, and 3 more variables: percent_pos <dbl>,
## #   IndivLevel <chr>, viruses <chr>
```

```
#write.csv(S1Table,'S1Table.csv')
```

##### Figure 4

###### Build out map side of figure

Get Vietnam map here from https://biogeo.ucdavis.edu/data/gadm3.6/Rsp/gadm36\_VNM\_1\_sp.rds

```
dataProjected<- readRDS(url("https://biogeo.ucdavis.edu/data/gadm3.6/Rsp/gadm36_VNM_1_sp.rds",method="libcurl"))
```

Add in color coding for each province

```
dataProjected@data$colorchoice <- rep("white", times=length(dataProjected@data$VARNAME_1))
```

```
## Loading required package: sp
```

```
dataProjected@data$colorchoice[dataProjected@data$VARNAME_1=="Dong Nai"] <-'grey'
dataProjected@data$colorchoice[dataProjected@data$VARNAME_1=="Dong Thap"] <-'grey'
dataProjected@data$colorchoice[dataProjected@data$VARNAME_1=="Soc Trang"] <-'grey'
```

Convert to sf object

```
dataProjected_sf <- st_as_sf(dataProjected)
```

Create data for labeling the 3 provinces

```
labeldata <- st_centroid(dataProjected_sf[dataProjected_sf$VARNAME_1 %in% c("Dong Nai","Dong Thap", "Soc Trang"),])
```

create map

```
ggViet <-ggplot(data=dataProjected_sf)+
  geom_sf(aes(fill=colorchoice), color='black',size =0.5) +
  scale_fill_identity("Province status",
    labels=c("Sampling","No sampling"),
    guide="legend")+
    geom_text_repel(data = labeldata,
                  aes(st_coordinates(labeldata)[,1],st_coordinates(labeldata)[,2],label = VARNAME_1),
                  #fontface = "bold",
                  nudge_x = c(-2, -2,  1),
                  nudge_y = c( 2,  1, -1),
                  size = 10*0.352777778) +  
  coord_sf() +
  theme(legend.text = element_text(size=10),
        legend.position = c(0.23,0.5),
        rect = element_blank(),
        axis.ticks = element_blank(),
        plot.margin=unit(c(0,0,0,0.5),"cm"),
        axis.text.y = element_blank(),
        axis.text.x = element_blank(),
        axis.title.y=element_blank(),
        axis.title.x=element_blank())


print(ggViet)
```

###### Build out panel side of figure

Create data and panel for each province. First, clarify taxomic group for the figure:

```
H1$plot_taxa <-as.character(H1$species_scientific_name)
H1$plot_taxa[H1$species_scientific_name=="Chiroptera"] <-'Microchiroptera'
H1$plot_taxa[H1$primary_interface_group=="trader"] <-'Field Rats'
H1$plot_taxa[H1$primary_interface_group=="large market"] <-'Field Rats'
H1$plot_taxa[H1$primary_interface_group=="restaurant"] <-'Field Rats'
H1$plot_taxa[H1$genus=="Pteropus"] <-"Pteropodidae"
H1$plot_taxa[H1$genus=="Cynopterus"] <-"Pteropodidae"
H1$plot_taxa[H1$genus=="Rhizomys"] <-"Rhizomys"
H1$plot_taxa <- as.factor(H1$plot_taxa)
H1$plot_taxa <- factor(H1$plot_taxa, levels = c("Field Rats", "Hystrix brachyura", "Rhizomys", "Rattus argentiventer", "Sciuridae", "Microchiroptera", "Pteropodidae"))
H1$primary_interface_group_2 <- H1$primary_interface_group 
levels(H1$primary_interface_group_2) <- c("trader", "large\nmarket", "restaurant", "natural\nroost", "bat\nguano\nfarm","wildlife\nfarm")
```

create summary data for figure

```
F4tab<- H1 %>%
  group_by(state_prov, primary_interface_group_2, plot_taxa) %>%
  summarize(ind_count=length(unique(animal_id)))
```

Bar chart panels

```
Fig4<-ggplot(F4tab, aes(x=primary_interface_group_2, y=ind_count, fill=plot_taxa)) +
      scale_fill_brewer(palette = "Set1") +
      geom_bar(stat='identity') +
      facet_wrap(~state_prov, ncol=1) +
      ylab("Individual count") +
      xlab("Interface") +
      labs(fill = "Taxa group") +
      theme_bw() + 
      theme(axis.text=element_text(size=8),
      #      axis.text.x = element_text(angle=45, hjust=1),
            axis.title = element_text(face="bold"), 
            legend.text =element_text(size=8), 
            legend.title =element_text(size=10),
            strip.text = element_text(size = 8),
            plot.margin=unit(c(0.25,0,0.25,0),"cm"))
print(Fig4)
```

combine map and bar charts for final Fig4:

```
grid.arrange(ggViet, Fig4, ncol = 3, 
             layout_matrix = cbind(c(1,1), c(2,2), c(2,2)))
```

##### Paragraph 2: Number of sites and proportion of positive sites

Number of sites

```
length(unique(H1$site_name))
```

```
## [1] 70
```

Number of sites where coronavirus was detected

```
length(unique(H1[H1$confirmation_result=="Positive","site_name"]))
```

```
## [1] 58
```

Compile number of sites with positive samples and site-level prevalence by interface and taxa\_group

```
H2.1i<- H1%>%
  group_by(taxa_group,primary_interface_group,site_name) %>%
  summarize(pos_site=sum(sum(confirmation_result=="Positive")>0))

H2.1i %>% 
  group_by(taxa_group,primary_interface_group) %>% 
  summarize(site_count=length(unique(site_name)),pos_sites=sum(pos_site),siteprev=pos_sites/site_count)
```

```
## # A tibble: 6 x 5
## # Groups:   taxa_group [2]
##   taxa_group       primary_interface_group site_count pos_sites siteprev
##   <fct>            <fct>                        <int>     <int>    <dbl>
## 1 Bats             natural roost                    1         1    1    
## 2 Bats             bat guano farm                  17        16    0.941
## 3 Rodents & Shrews trader                           8         8    1    
## 4 Rodents & Shrews large market                    14        14    1    
## 5 Rodents & Shrews restaurant                       2         2    1    
## 6 Rodents & Shrews wildlife farm                   28        17    0.607
```

##### Table 1

This code is producing different sections of Table 1

Main structure:

```
T1<- H1 %>%
  group_by(taxa_group, primary_interface_group, new_genus) %>%
  summarize(ind_count=length(unique(animal_id)), 
            ind_count_pos=length(unique(animal_id[confirmation_result=="Positive"])),
            percent_pos = ind_count_pos/ind_count*100,
            site_count=length(unique(site_name)),
            pos_sites=length(unique(site_name[confirmation_result=="Positive"])),
            siteprev=pos_sites/site_count*100,
            SiteLevel= paste0(round(siteprev,digits = 1), "% (",pos_sites,"/",site_count,")"),
            IndivLevel= paste0(round(percent_pos,digits = 1), "% (",ind_count_pos,"/",ind_count,")"),
            viruses = paste0(unique(virus_ind), collapse=", "))%>% 
  select(-c(4:9)) %>% 
  arrange(desc(taxa_group))
T1
```

```
## # A tibble: 10 x 6
## # Groups:   taxa_group, primary_interface_group [6]
##    taxa_group  primary_interfac… new_genus  SiteLevel  IndivLevel viruses       
##    <fct>       <fct>             <chr>      <chr>      <chr>      <chr>         
##  1 Rodents & … trader            Field Rats 100% (8/8) 20.7% (39… Murine corona…
##  2 Rodents & … large market      Field Rats 100% (14/… 32% (116/… Murine corona…
##  3 Rodents & … restaurant        Field Rats 100% (2/2) 55.6% (84… Murine corona…
##  4 Rodents & … wildlife farm     Hystrix    47.8% (11… 6% (20/33… Bat coronavir…
##  5 Rodents & … wildlife farm     Rattus     100% (1/1) 100% (1/1) Bat coronavir…
##  6 Rodents & … wildlife farm     Rhizomys   45.5% (5/… 6.2% (6/9… Infectious br…
##  7 Rodents & … wildlife farm     Sciuridae  0% (0/1)   0% (0/1)   NA            
##  8 Bats        natural roost     Cynopterus 0% (0/1)   0% (0/2)   NA            
##  9 Bats        natural roost     Pteropus   100% (1/1) 6.7% (4/6… PREDICT_CoV-1…
## 10 Bats        bat guano farm    Microchir… 94.1% (16… 74.8% (23… Bat coronavir…
```

Number of individuals found with each virus:

```
T1v<- H1 %>%
  group_by(taxa_group, primary_interface_group, new_genus, virus_group) %>%
  summarize(ind_count_virus=length(unique(animal_id))) %>% 
 mutate(virus_ind_count=paste0(virus_group, " (", ind_count_virus,")"))%>% 
  group_by(taxa_group, primary_interface_group, new_genus) %>% 
  summarize(viral_species=paste0(virus_ind_count ,collapse = ', ')) %>% 
    arrange(desc(taxa_group)) 
T1v
```

```
## # A tibble: 10 x 4
## # Groups:   taxa_group, primary_interface_group [6]
##    taxa_group   primary_interface_… new_genus  viral_species                    
##    <fct>        <fct>               <chr>      <chr>                            
##  1 Rodents & S… trader              Field Rats Longquan Aa mouse coronavirus (5…
##  2 Rodents & S… large market        Field Rats Longquan Aa mouse coronavirus (3…
##  3 Rodents & S… restaurant          Field Rats Longquan Aa mouse coronavirus (2…
##  4 Rodents & S… wildlife farm       Hystrix    Bat coronavirus 512/2005 (19), I…
##  5 Rodents & S… wildlife farm       Rattus     Bat coronavirus 512/2005 (1), NU…
##  6 Rodents & S… wildlife farm       Rhizomys   Bat coronavirus 512/2005 (5), In…
##  7 Rodents & S… wildlife farm       Sciuridae  NULL (1)                         
##  8 Bats         natural roost       Cynopterus NULL (2)                         
##  9 Bats         natural roost       Pteropus   NULL (59), PREDICT_CoV-17 (3), P…
## 10 Bats         bat guano farm      Microchir… Bat coronavirus 512/2005 (216), …
```

Coinfections column

```
coinfects<-H1[grep('\\|',H1$virus_ind),]

T1coinf<-coinfects %>%
  group_by(taxa_group, primary_interface_group, new_genus) %>%
  summarize(ind_count=length(unique(animal_id))) %>% 
  arrange(desc(taxa_group))
T1coinf
```

```
## # A tibble: 4 x 4
## # Groups:   taxa_group, primary_interface_group [4]
##   taxa_group       primary_interface_group new_genus       ind_count
##   <fct>            <fct>                   <chr>               <int>
## 1 Rodents & Shrews trader                  Field Rats              2
## 2 Rodents & Shrews large market            Field Rats             18
## 3 Rodents & Shrews restaurant              Field Rats              6
## 4 Bats             bat guano farm          Microchiroptera        21
```

##### Paragraph 3 Field rodents

Subset the data for rodents:

```
H4<-subset(H1, taxa_group=="Rodents & Shrews")
length(unique(H4$animal_id))
```

```
## [1] 1131
```

Proportion positive at site and animal level by province:

```
 H4 %>%
  filter(primary_interface_group!="wildlife farm") %>% 
  group_by(state_prov) %>%
  summarize(site_count=length(unique(site_name)),pos_sites=length(unique(site_name[confirmation_result=="Positive"])),siteprev=pos_sites/site_count*100,
            ind_count=length(unique(animal_id)), 
            ind_count_pos=length(unique(animal_id[confirmation_result=="Positive"])),
            percent_pos = ind_count_pos/ind_count*100) %>% 
  rowwise() %>% 
  mutate(confint=paste0(round(100*prop.test(ind_count_pos,ind_count, conf.level=0.95)$conf.int,1),collapse = ' - '))
```

```
## Source: local data frame [2 x 8]
## Groups: <by row>
## 
## # A tibble: 2 x 8
##   state_prov site_count pos_sites siteprev ind_count ind_count_pos percent_pos
##   <fct>           <int>     <int>    <dbl>     <int>         <int>       <dbl>
## 1 Dong Thap          16        16      100       373           129        34.6
## 2 Soc Trang           8         8      100       329           110        33.4
## # … with 1 more variable: confint <chr>
```

Overall proportion of positive field rats:

```
H4 %>%
  filter(primary_interface_group!="wildlife farm") %>% 
  summarize(site_count=length(unique(site_name)),pos_sites=length(unique(site_name[confirmation_result=="Positive"])),siteprev=pos_sites/site_count*100,
            ind_count=length(unique(animal_id)), 
            ind_count_pos=length(unique(animal_id[confirmation_result=="Positive"])),
            percent_pos = ind_count_pos/ind_count*100)  %>% 
  rowwise() %>% 
  mutate(confint=paste0(round(100*prop.test(ind_count_pos,ind_count, conf.level=0.95)$conf.int,1),collapse = ' - '))
```

```
## Source: local data frame [1 x 7]
## Groups: <by row>
## 
## # A tibble: 1 x 7
##   site_count pos_sites siteprev ind_count ind_count_pos percent_pos confint    
##        <int>     <int>    <dbl>     <int>         <int>       <dbl> <chr>      
## 1         24        24      100       702           239        34.0 30.6 - 37.7
```

proportion of positive field rats by site:

```
H4bysite <- H4 %>%
  filter(primary_interface_group!="wildlife farm") %>% 
  group_by(site_name) %>%
  summarize(site_count=length(unique(site_name)),pos_sites=length(unique(site_name[confirmation_result=="Positive"])),siteprev=pos_sites/site_count*100,
            ind_count=length(unique(animal_id)), 
            ind_count_pos=length(unique(animal_id[confirmation_result=="Positive"])),
            percent_pos = ind_count_pos/ind_count*100) %>% 
  rowwise() %>% 
  mutate(confint=paste0(round(100*prop.test(ind_count_pos,ind_count, conf.level=0.95)$conf.int,1),collapse = ' - '))

H4bysite
```

```
## Source: local data frame [24 x 8]
## Groups: <by row>
## 
## # A tibble: 24 x 8
##    site_name site_count pos_sites siteprev ind_count ind_count_pos percent_pos
##    <fct>          <int>     <int>    <dbl>     <int>         <int>       <dbl>
##  1 Rat mark…          1         1      100         9             5       55.6 
##  2 Rat mark…          1         1      100        20             9       45   
##  3 Rat mark…          1         1      100        15             5       33.3 
##  4 Rat mark…          1         1      100        20             5       25   
##  5 Rat mark…          1         1      100        26             6       23.1 
##  6 Rat mark…          1         1      100        26             2        7.69
##  7 Rat mark…          1         1      100        20             7       35   
##  8 Rat mark…          1         1      100        22            11       50   
##  9 Rat mark…          1         1      100        18             4       22.2 
## 10 Rat mark…          1         1      100        19             6       31.6 
## # … with 14 more rows, and 1 more variable: confint <chr>
```

```
H4bysite%>% 
summarize(max_site=max(percent_pos),min_site=min(percent_pos))
```

```
## # A tibble: 24 x 2
##    max_site min_site
##       <dbl>    <dbl>
##  1    55.6     55.6 
##  2    45       45   
##  3    33.3     33.3 
##  4    25       25   
##  5    23.1     23.1 
##  6     7.69     7.69
##  7    35       35   
##  8    50       50   
##  9    22.2     22.2 
## 10    31.6     31.6 
## # … with 14 more rows
```

Proportion of positive rodents by sub-interface type

```
 H4 %>%
  filter(primary_interface_group!="wildlife farm") %>% 
  group_by(primary_interface_group) %>%
  summarize(site_count=length(unique(site_name)),pos_sites=length(unique(site_name[confirmation_result=="Positive"])),siteprev=pos_sites/site_count*100,
            ind_count=length(unique(animal_id)), 
            ind_count_pos=length(unique(animal_id[confirmation_result=="Positive"])),
            percent_pos = ind_count_pos/ind_count*100)%>% 
  rowwise() %>% 
  mutate(confint=paste0(round(100*prop.test(ind_count_pos,ind_count, conf.level=0.95)$conf.int,1),collapse = ' - '))
```

```
## Source: local data frame [3 x 8]
## Groups: <by row>
## 
## # A tibble: 3 x 8
##   primary_interfa… site_count pos_sites siteprev ind_count ind_count_pos
##   <fct>                 <int>     <int>    <dbl>     <int>         <int>
## 1 trader                    8         8      100       188            39
## 2 large market             14        14      100       363           116
## 3 restaurant                2         2      100       151            84
## # … with 2 more variables: percent_pos <dbl>, confint <chr>
```

Proportion of positive rodents by season

```
 H4 %>%
  filter(primary_interface_group!="wildlife farm") %>% 
  group_by(hseason) %>%
  summarize(site_count=length(unique(site_name)),pos_sites=length(unique(site_name[confirmation_result=="Positive"])),
            siteprev=pos_sites/site_count*100,
            ind_count=length(unique(animal_id)), 
            ind_count_pos=length(unique(animal_id[confirmation_result=="Positive"])),
            percent_pos = ind_count_pos/ind_count*100)%>% 
  rowwise() %>% 
  mutate(confint=paste0(round(100*prop.test(ind_count_pos,ind_count, conf.level=0.95)$conf.int,1),collapse = ' - '))
```

```
## Source: local data frame [2 x 8]
## Groups: <by row>
## 
## # A tibble: 2 x 8
##   hseason site_count pos_sites siteprev ind_count ind_count_pos percent_pos
##   <chr>        <int>     <int>    <dbl>     <int>         <int>       <dbl>
## 1 Dry              2         2      100       130            29        22.3
## 2 Wet             22        22      100       572           210        36.7
## # … with 1 more variable: confint <chr>
```

###### Testing effect of season and sub-interface on risk of infection of rodent along supply chain

Setting up the data for uni- and multi-variable models

```
H1st<-H1 %>% 
  filter(state_prov %in% c("Dong Thap", "Soc Trang") & primary_interface_group %in% c("trader","large market","restaurant")) %>% 
  group_by(new_genus,animal_id) %>%
  summarize(coronavirus=length(unique(animal_id[confirmation_result=="Positive"])),
            season = unique(hseason),
            site_name = unique(site_name),
            interface= unique(primary_interface_group))


H1st$season <- as.factor(H1st$season)
H1st$season <- factor(H1st$season, levels = c("Dry", "Wet"))
H1st$interface<- as.factor(H1st$interface)
H1st$interface<- factor(H1st$interface, levels = c("trader","large market","restaurant"))
H1st
```

```
## # A tibble: 702 x 6
## # Groups:   new_genus [1]
##    new_genus  animal_id coronavirus season site_name                  interface 
##    <chr>      <fct>           <int> <fct>  <fct>                      <fct>     
##  1 Field Rats VN13M0001           0 Dry    Restaurant - Soc Trang     restaurant
##  2 Field Rats VN13M0002           1 Dry    Restaurant - Soc Trang     restaurant
##  3 Field Rats VN13M0003           1 Dry    Restaurant - Soc Trang     restaurant
##  4 Field Rats VN13M0004           0 Dry    Wet market (Whole sale se… large mar…
##  5 Field Rats VN13M0005           1 Dry    Restaurant - Soc Trang     restaurant
##  6 Field Rats VN13M0006           0 Dry    Restaurant - Soc Trang     restaurant
##  7 Field Rats VN13M0007           1 Dry    Restaurant - Soc Trang     restaurant
##  8 Field Rats VN13M0008           1 Dry    Restaurant - Soc Trang     restaurant
##  9 Field Rats VN13M0009           1 Dry    Restaurant - Soc Trang     restaurant
## 10 Field Rats VN13M0010           0 Dry    Wet market (Whole sale se… large mar…
## # … with 692 more rows
```

Univariate models for season and interface:

```
m_st.1<-glm(coronavirus~season, family = binomial, data = H1st)
summary(m_st.1)
```

```
## 
## Call:
## glm(formula = coronavirus ~ season, family = binomial, data = H1st)
## 
## Deviance Residuals: 
##     Min       1Q   Median       3Q      Max  
## -0.9566  -0.9566  -0.9566   1.4157   1.7322  
## 
## Coefficients:
##             Estimate Std. Error z value Pr(>|z|)    
## (Intercept)  -1.2478     0.2107  -5.923 3.16e-09 ***
## seasonWet     0.7033     0.2278   3.087  0.00202 ** 
## ---
## Signif. codes:  0 '***' 0.001 '**' 0.01 '*' 0.05 '.' 0.1 ' ' 1
## 
## (Dispersion parameter for binomial family taken to be 1)
## 
##     Null deviance: 900.44  on 701  degrees of freedom
## Residual deviance: 890.08  on 700  degrees of freedom
## AIC: 894.08
## 
## Number of Fisher Scoring iterations: 4
```

```
exp(cbind(OR = coef(m_st.1), confint(m_st.1)))
```

```
## Waiting for profiling to be done...
```

```
##                    OR     2.5 %    97.5 %
## (Intercept) 0.2871287 0.1867155 0.4277976
## seasonWet   2.0203848 1.3082077 3.2047957
```

```
m_st.2<-glm(coronavirus~interface, family = binomial, data = H1st)
summary(m_st.2)
```

```
## 
## Call:
## glm(formula = coronavirus ~ interface, family = binomial, data = H1st)
## 
## Deviance Residuals: 
##     Min       1Q   Median       3Q      Max  
## -1.2748  -0.8775  -0.6819   1.0830   1.7736  
## 
## Coefficients:
##                       Estimate Std. Error z value Pr(>|z|)    
## (Intercept)            -1.3404     0.1799  -7.452 9.19e-14 ***
## interfacelarge market   0.5846     0.2122   2.755  0.00587 ** 
## interfacerestaurant     1.5665     0.2433   6.439 1.20e-10 ***
## ---
## Signif. codes:  0 '***' 0.001 '**' 0.01 '*' 0.05 '.' 0.1 ' ' 1
## 
## (Dispersion parameter for binomial family taken to be 1)
## 
##     Null deviance: 900.44  on 701  degrees of freedom
## Residual deviance: 854.25  on 699  degrees of freedom
## AIC: 860.25
## 
## Number of Fisher Scoring iterations: 4
```

```
exp(cbind(OR = coef(m_st.2), confint(m_st.2)))
```

```
## Waiting for profiling to be done...
```

```
##                             OR    2.5 %    97.5 %
## (Intercept)           0.261745 0.181572 0.3682482
## interfacelarge market 1.794249 1.192172 2.7436193
## interfacerestaurant   4.789897 2.993666 7.7802354
```

Multivariable model:

```
m_st.4<-glmer(coronavirus~interface+season +(1|site_name), family = binomial, data = H1st)
summary(m_st.4)
```

```
## Generalized linear mixed model fit by maximum likelihood (Laplace
##   Approximation) [glmerMod]
##  Family: binomial  ( logit )
## Formula: coronavirus ~ interface + season + (1 | site_name)
##    Data: H1st
## 
##      AIC      BIC   logLik deviance df.resid 
##    801.6    824.4   -395.8    791.6      697 
## 
## Scaled residuals: 
##     Min      1Q  Median      3Q     Max 
## -1.4641 -0.6956 -0.3648  0.7019  3.1641 
## 
## Random effects:
##  Groups    Name        Variance Std.Dev.
##  site_name (Intercept) 0.4105   0.6407  
## Number of obs: 702, groups:  site_name, 24
## 
## Fixed effects:
##                       Estimate Std. Error z value Pr(>|z|)    
## (Intercept)            -2.9838     0.6842  -4.361 1.29e-05 ***
## interfacelarge market   0.7892     0.3693   2.137 0.032567 *  
## interfacerestaurant     2.3052     0.6441   3.579 0.000345 ***
## seasonWet               1.5927     0.6165   2.583 0.009788 ** 
## ---
## Signif. codes:  0 '***' 0.001 '**' 0.01 '*' 0.05 '.' 0.1 ' ' 1
## 
## Correlation of Fixed Effects:
##             (Intr) intrfm intrfc
## intrfclrgmr -0.466              
## intrfcrstrn -0.622  0.430       
## seasonWet   -0.902  0.134  0.469
```

check if any site was sampled more than once

```
H1 %>% 
  filter(state_prov %in% c("Dong Thap", "Soc Trang") & primary_interface_group %in% c("trader","large market","restaurant")) %>% 
  group_by(site_name) %>% 
  summarize(n_visit=length(unique(event_name)))
```

```
## # A tibble: 24 x 2
##    site_name                   n_visit
##    <fct>                         <int>
##  1 Rat market CLC1 - Dong Thap       1
##  2 Rat market CLC2 - Dong Thap       1
##  3 Rat market CLC3 - Dong Thap       1
##  4 Rat market CLD - Dong Thap        1
##  5 Rat market CT - Dong Thap         1
##  6 Rat market HND - Dong Thap        1
##  7 Rat market HNT - Dong Thap        1
##  8 Rat market LVO - Dong Thap        1
##  9 Rat market LVU - Dong Thap        1
## 10 Rat market MX1 - Soc Trang        1
## # … with 14 more rows
```

Site only sampled once, so practically, both season and interface are site-level variables, and confounding effects with other site-level characteristics cannot be excluded. Although the GLMM is the right modeling approach for this data structure, caution required in the interpretation due to sampling design limitations

##### Table S2 Field rodent multi-variable model with site as a random effect

```
TS2 <- as.data.frame(exp(confint(m_st.4)))
```

```
## Computing profile confidence intervals ...
```

```
TS2$OR <- NA
TS2$OR[3:5] <- exp(summary(m_st.4)$coefficients[2:4,1])
TS2[3:5,c(3,1,2)]
```

```
##                              OR    2.5 %    97.5 %
## interfacelarge market  2.201718 1.045097  4.751261
## interfacerestaurant   10.025683 2.665258 39.453799
## seasonWet              4.916909 1.375610 18.024858
```

##### Figure 5 Graph representation of these results -

```
RodT_int <- H1 %>% 
  filter(state_prov %in% c("Dong Thap", "Soc Trang", "Dong Nai") & taxa_group=="Rodents & Shrews" & primary_interface_group %in% c("large market", "restaurant", "trader")) %>% 
   group_by(primary_interface_group) %>%
   summarize(N=length(unique(animal_id)), 
            Positive=length(unique(animal_id[confirmation_result=="Positive"])),
            Proportion = 100*Positive/N,
            lab = paste0(round(Proportion,1), "%\n n=", N),
            L_CI = round(100*prop.test(Positive,N, conf.level=0.95)$conf.int[1],1),
            U_CI = round(100*prop.test(Positive,N, conf.level=0.95)$conf.int[2],1)) %>% 
  as.data.frame()

Fig5 <- ggplot(RodT_int,aes(primary_interface_group, Proportion)) +
  geom_col(fill="lightgrey") + 
  geom_errorbar(aes(ymin = ifelse(L_CI<0,0,L_CI), ymax = U_CI), width = .5, color="grey", alpha=0.9) +
  ggtitle("Proportion of coronavirus-positive field rats") + 
  xlab("Field rat supply chain interface") +
  geom_text(aes(label = lab, hjust = .5, vjust= 0.50), position = position_dodge(width = 0), 
            angle = 0)+
    theme_bw()
Fig5
```

```
#tiff("Fig 5.tif", height = 4, width = 4, units = 'in', res=300)
#grid.arrange(Fig5, ncol = 1, layout_matrix = cbind(c(1,1)))
#dev.off()
```

##### Paragraph 4 - Probability of detection with different sample types among positive field rats with more than one sample

Select animals id from the rodent trade that:

- have multiple sample types
- are positive

```
Rp<-H4 %>% 
  filter(confirmation_result=="Positive")

Rmsp<-H4 %>% 
    filter(primary_interface_group!="wildlife farm") %>% 
  group_by(animal_id) %>% 
  summarize(spl_type=length(unique(specimen_type))) %>% 
  filter(spl_type>1)

H5<-H4 %>% 
  filter(animal_id %in% Rmsp$animal_id & animal_id %in% unique(Rp$animal_id)) 
length(unique(H5$animal_id))
```

```
## [1] 220
```

then look at proportion of positive by sample types

```
H5%>% 
  group_by(specimen_type) %>% 
  summarize(ind_count=length(unique(animal_id)), 
            ind_count_pos=length(unique(animal_id[confirmation_result=="Positive"])),
            percent_pos = round(ind_count_pos/ind_count*100,1)) %>% 
  rowwise() %>% 
  mutate(confint=paste0(round(100*prop.test(ind_count_pos,ind_count, conf.level=0.95)$conf.int,1),collapse = ' - '))
```

```
## Source: local data frame [8 x 5]
## Groups: <by row>
## 
## # A tibble: 8 x 5
##   specimen_type         ind_count ind_count_pos percent_pos confint    
##   <fct>                     <int>         <int>       <dbl> <chr>      
## 1 brain                        16             5        31.2 12.1 - 58.5
## 2 feces                         2             1        50   9.5 - 90.5 
## 3 kidney                       13             3        23.1 6.2 - 54   
## 4 lung                         51            27        52.9 38.6 - 66.8
## 5 oral swab                   219           175        79.9 73.9 - 84.9
## 6 small intestine             155            80        51.6 43.5 - 59.7
## 7 spleen                        1             1       100   5.5 - 100  
## 8 urine/urogenital swab         1             0         0   0 - 94.5
```

##### Paragraph 5 rodent farm results

Proportion of positive animals by genus in wildife farms

```
H4 %>%
    filter(primary_interface_group=="wildlife farm") %>% 
  group_by(new_genus) %>%
  summarize(ind_count=length(unique(animal_id)), 
            ind_count_pos=length(unique(animal_id[confirmation_result=="Positive"])),
            percent_pos = ind_count_pos/ind_count*100) %>% 
  rowwise() %>% 
  mutate(confint=paste0(round(100*prop.test(ind_count_pos,ind_count, conf.level=0.95)$conf.int,1),collapse = ' - '))
```

```
## Source: local data frame [4 x 5]
## Groups: <by row>
## 
## # A tibble: 4 x 5
##   new_genus ind_count ind_count_pos percent_pos confint   
##   <chr>         <int>         <int>       <dbl> <chr>     
## 1 Hystrix         331            20        6.04 3.8 - 9.3 
## 2 Rattus            1             1      100    5.5 - 100 
## 3 Rhizomys         96             6        6.25 2.6 - 13.6
## 4 Sciuridae         1             0        0    0 - 94.5
```

overall, at site- and animal-level

```
H4 %>%
  filter(primary_interface_group=="wildlife farm") %>% 
  group_by(state_prov) %>%
  summarize(site_count=length(unique(site_name)),pos_sites=length(unique(site_name[confirmation_result=="Positive"])),siteprev=pos_sites/site_count*100,
            ind_count=length(unique(animal_id)), 
            ind_count_pos=length(unique(animal_id[confirmation_result=="Positive"])),
            percent_pos = ind_count_pos/ind_count*100) %>% 
  rowwise() %>% 
  mutate(confint=paste0(round(100*prop.test(ind_count_pos,ind_count, conf.level=0.95)$conf.int,1),collapse = ' - '))
```

```
## Source: local data frame [1 x 8]
## Groups: <by row>
## 
## # A tibble: 1 x 8
##   state_prov site_count pos_sites siteprev ind_count ind_count_pos percent_pos
##   <fct>           <int>     <int>    <dbl>     <int>         <int>       <dbl>
## 1 Dong Nai           28        17     60.7       429            27        6.29
## # … with 1 more variable: confint <chr>
```

and disaggregated by genus and season (showing limitations in sampling design):

```
H4 %>%
    filter(primary_interface_group=="wildlife farm") %>% 
  group_by(new_genus, hseason) %>%
  summarize(ind_count=length(unique(animal_id)), 
            ind_count_pos=length(unique(animal_id[confirmation_result=="Positive"])),
            percent_pos = ind_count_pos/ind_count*100)
```

```
## # A tibble: 6 x 5
## # Groups:   new_genus [4]
##   new_genus hseason ind_count ind_count_pos percent_pos
##   <chr>     <chr>       <int>         <int>       <dbl>
## 1 Hystrix   Dry           296            20        6.76
## 2 Hystrix   Wet            35             0        0   
## 3 Rattus    Wet             1             1      100   
## 4 Rhizomys  Dry            95             6        6.32
## 5 Rhizomys  Wet             1             0        0   
## 6 Sciuridae Dry             1             0        0
```

and season only (aggregated on genus):

```
H4 %>%
    filter(primary_interface_group=="wildlife farm") %>% 
  group_by(hseason) %>%
  summarize(ind_count=length(unique(animal_id)), 
            ind_count_pos=length(unique(animal_id[confirmation_result=="Positive"])),
            percent_pos = ind_count_pos/ind_count*100)
```

```
## # A tibble: 2 x 4
##   hseason ind_count ind_count_pos percent_pos
##   <chr>       <int>         <int>       <dbl>
## 1 Dry           392            26        6.63
## 2 Wet            37             1        2.70
```

##### Paragraph 6 Bat findings

Subset the data for bats

```
H3<-subset(H1, taxa_group=="Bats")
length(unique(H3$animal_id)) #check
```

```
## [1] 375
```

Look at proportion positive per taxa based on animal ID

```
H3.1 <- H3 %>%
  group_by(taxa_group, new_genus) %>%
  summarize(ind_count=length(unique(animal_id)),
            ind_count_pos=length(unique(animal_id[confirmation_result=="Positive"])),
            ind_percent_pos = ind_count_pos/ind_count*100,
            ) %>% 
  rowwise() %>% 
  mutate(confint=paste0(round(100*prop.test(ind_count_pos,ind_count, conf.level=0.95)$conf.int,1),collapse = ' - '))
H3.1
```

```
## Source: local data frame [3 x 6]
## Groups: <by row>
## 
## # A tibble: 3 x 6
##   taxa_group new_genus       ind_count ind_count_pos ind_percent_pos confint    
##   <fct>      <chr>               <int>         <int>           <dbl> <chr>      
## 1 Bats       Cynopterus              2             0            0    0 - 80.2   
## 2 Bats       Microchiroptera       313           234           74.8  69.5 - 79.4
## 3 Bats       Pteropus               60             4            6.67 2.2 - 17
```

###### Seasonality for the Soc Trang bat pagoda

Look at proportion positive per season

```
H3.2 <- H3 %>%
  filter(new_genus=="Pteropus") %>% 
  group_by(hseason) %>%
  summarize(ind_count=length(unique(animal_id)),
            ind_count_pos=length(unique(animal_id[confirmation_result=="Positive"])),
            ind_percent_pos = ind_count_pos/ind_count*100,
            ) %>% 
  rowwise() %>% 
  mutate(confint=paste0(round(100*prop.test(ind_count_pos,ind_count, conf.level=0.95)$conf.int,1),collapse = ' - '))
H3.2
```

```
## Source: local data frame [2 x 5]
## Groups: <by row>
## 
## # A tibble: 2 x 5
##   hseason ind_count ind_count_pos ind_percent_pos confint   
##   <chr>       <int>         <int>           <dbl> <chr>     
## 1 Dry            49             1            2.04 0.1 - 12.2
## 2 Wet            11             3           27.3  7.3 - 60.7
```

Fisher exact test:

```
H1bp<-H1 %>% 
  filter(site_name=="Bat pagoda - Soc Trang", genus=="Pteropus") %>% 
  group_by(animal_id) %>%
  summarize(coronavirus=length(unique(animal_id[confirmation_result=="Positive"])),
            season = unique(hseason),
            site_name = unique(site_name))
fisher.test(table(H1bp$season,H1bp$coronavirus))
```

```
## 
##  Fisher's Exact Test for Count Data
## 
## data:  table(H1bp$season, H1bp$coronavirus)
## p-value = 0.01726
## alternative hypothesis: true odds ratio is not equal to 1
## 95 percent confidence interval:
##    1.178107 956.801500
## sample estimates:
## odds ratio 
##   16.63662
```

This suggest an increased risk of being positive for cov during wet season for the Soc Trang pteropus

#### Phylogenetic analysis

List viral taxonomic units

```
H1$virus[H1$virus=="NULL"] <- NA
H1 %>% 
  distinct(virus)
```

```
##                                         virus
## 1          strain of Bat coronavirus 512/2005
## 2                              PREDICT_CoV-35
## 3                                        <NA>
## 4                              PREDICT_CoV-17
## 5                strain of Murine coronavirus
## 6     strain of Longquan Aa mouse coronavirus
## 7 strain of Infectious bronchitis virus (IBV)
```

Number of viruses obtained with Watanbe protocol:

```
H1 %>% 
  filter(test_requested_protocol=='Modified Watanabe et al, RdRp gene') %>% 
  distinct(virus)
```

```
##                                     virus
## 1      strain of Bat coronavirus 512/2005
## 2                                    <NA>
## 3                          PREDICT_CoV-35
## 4                          PREDICT_CoV-17
## 5            strain of Murine coronavirus
## 6 strain of Longquan Aa mouse coronavirus
```

Number of viruses obtained with Quan protocol:

```
H1 %>% 
  filter( test_requested_protocol=='Quan et al, RdRp gene')%>% 
  distinct(virus)
```

```
##                                         virus
## 1          strain of Bat coronavirus 512/2005
## 2                              PREDICT_CoV-35
## 3                                        <NA>
## 4                strain of Murine coronavirus
## 5     strain of Longquan Aa mouse coronavirus
## 6                              PREDICT_CoV-17
## 7 strain of Infectious bronchitis virus (IBV)
```

```
allpos<-H1 %>%
  filter(confirmation_result=="Positive") %>% 
  group_by(animal_id) %>%
  summarize(ind_count=length(unique(animal_id)),
            want_pos=length(unique(animal_id[test_requested_protocol=="Modified Watanabe et al, RdRp gene" & confirmation_result=="Positive"])),
            quan_pos=length(unique(animal_id[test_requested_protocol=="Quan et al, RdRp gene" & confirmation_result=="Positive"])),
            both_tests_pos = ifelse(want_pos+quan_pos==2,1,0))

colSums(allpos[,2:5])
```

```
##      ind_count       want_pos       quan_pos both_tests_pos 
##            504            433            410            339
```

Viruses found in bats

```
H1 %>% 
  filter(virus_group=="PREDICT_CoV-17") %>% 
  group_by(new_genus) %>% 
  summarize(n_Infected_with_PCov17=length(unique(animal_id)))
```

```
## # A tibble: 2 x 2
##   new_genus       n_Infected_with_PCov17
##   <chr>                            <int>
## 1 Microchiroptera                      1
## 2 Pteropus                             3
```

```
H1 %>% 
  filter(virus_group=="PREDICT_CoV-35") %>% 
  group_by(new_genus) %>% 
  summarize(n_Infected_with_PCov35=length(unique(animal_id)))
```

```
## # A tibble: 2 x 2
##   new_genus       n_Infected_with_PCov35
##   <chr>                            <int>
## 1 Microchiroptera                     38
## 2 Pteropus                             1
```

```
H1 %>% 
  filter(virus_group=="Bat coronavirus 512/2005") %>% 
  group_by(new_genus) %>% 
  summarize(n_Infected_with_BCov5122005=length(unique(animal_id)))
```

```
## # A tibble: 4 x 2
##   new_genus       n_Infected_with_BCov5122005
##   <chr>                                 <int>
## 1 Hystrix                                  19
## 2 Microchiroptera                         216
## 3 Rattus                                    1
## 4 Rhizomys                                  5
```

See also Table 1.

Viruses found in rodents

```
H1 %>%
  group_by(taxa_group, new_genus, virus_group) %>%
  summarize(ind_count_virus=length(unique(animal_id))) %>% 
 mutate(virus_ind_count=paste0(virus_group, " (", ind_count_virus,")"))%>% 
  group_by(taxa_group, new_genus) %>% 
  summarize(viral_species=paste0(virus_ind_count ,collapse = ', ')) %>% 
    arrange(desc(taxa_group))
```

```
## # A tibble: 8 x 3
## # Groups:   taxa_group [2]
##   taxa_group     new_genus    viral_species                                     
##   <fct>          <chr>        <chr>                                             
## 1 Rodents & Shr… Field Rats   Longquan Aa mouse coronavirus (56), Murine corona…
## 2 Rodents & Shr… Hystrix      Bat coronavirus 512/2005 (19), Infectious bronchi…
## 3 Rodents & Shr… Rattus       Bat coronavirus 512/2005 (1), NULL (1)            
## 4 Rodents & Shr… Rhizomys     Bat coronavirus 512/2005 (5), Infectious bronchit…
## 5 Rodents & Shr… Sciuridae    NULL (1)                                          
## 6 Bats           Cynopterus   NULL (2)                                          
## 7 Bats           Microchirop… Bat coronavirus 512/2005 (216), NULL (126), PREDI…
## 8 Bats           Pteropus     NULL (59), PREDICT_CoV-17 (3), PREDICT_CoV-35 (1)
```

and see code for Table 1 above

Details on IBV detection:

```
H1 %>%
  filter(virus_group=="Infectious bronchitis virus (IBV)") %>% 
  group_by(primary_interface_group,taxa_group,species_scientific_name, site_name, id_certainty) %>%
  summarize(ind_count=length(unique(animal_id)))
```

```
## # A tibble: 2 x 6
## # Groups:   primary_interface_group, taxa_group, species_scientific_name,
## #   site_name [2]
##   primary_interfac… taxa_group species_scienti… site_name id_certainty ind_count
##   <fct>             <fct>      <fct>            <fct>     <fct>            <int>
## 1 wildlife farm     Rodents &… Hystrix brachyu… Farm 25 … field ID ce…         1
## 2 wildlife farm     Rodents &… Rhizomys sp.     Farm 4 -… field ID ce…         1
```
