## Supplementary material for "Coronavirus testing indicates transmission risk increases along wildlife supply chains for human consumption in Viet Nam, 2013-2014": S1 Table

**S1 Table. Summary of all testing results by genus, interface, sub-interface, sample types, sites, percentage of samples testing positive, and viral species.**

| Genus | Interface | Sub-interface | Sample type (sites) | % individual positive | Viral species (X\|X indicates co-infection) |
|---|---|---|---|---|---|
| *Cynopterus* | Human dwelling | Natural bat roost | Oral swab (1) | 0% (0/1) | |
| | | | Rectal swab (1) | 0% (0/2) | |
| *Pteropus* | Human dwelling | Natural bat roost | Feces (1) | 8.9% (4/45) | PREDICT_CoV-17, PREDICT_CoV-35 |
| | | | Oral swab (1) | 0% (0/13) | |
| | | | Rectal swab (1) | 0% (0/15) | |
| | | | Urine (1) | 0% (0/2) | |
| *Micro-chiroptera*[a] | Human dwelling | Bat guano farm | Feces (17) | 76.5% (234/306) | Bat coronavirus 512/2005, PREDICT_CoV-35, PREDICT_CoV-35\|Bat coronavirus 512/2005, PREDICT_CoV-17\|Bat coronavirus 512/2005 |
| | | Bat guano farm | Urine (2) | 0% (0/7) | |
| Field rats *(Rattus + Bandicota)* | Live rodent trade | Large market | Brain (5) | 8.9% (4/45) | Murine coronavirus |
| | | | Feces (1) | 0% (0/13) | |
| | | | Kidney (8) | 4.9% (3/61) | Murine coronavirus |
| | | | Lung (3) | 7.8% (4/51) | Murine coronavirus, Longquan aa coronavirus |
| | | | Oral swab (14) | 32.1% (88/274) | Murine coronavirus, Longquan aa coronavirus, Murine coronavirus \| Longquan aa coronavirus |
| | | | Rectal swab (1) | 0% (0/1) | |
| | | | Small intestine (13) | 22.6% (50/221) | Murine coronavirus, Longquan aa coronavirus, |

| | | | | | |
|---|---|---|---|---|---|
| | | | | | Murine coronavirus \| Longquan aa coronavirus |
| | | | Urine swab (1) | 0% (0/6) | |
| | | Restaurant | Feces (1) | 50.0% (1/2) | Longquan Aa mouse coronavirus |
| | | | Lung (2) | 27.1% (23/85) | Murine coronavirus, Longquan aa coronavirus |
| | | | Oral swab (2) | 51.3% (61/119) | Murine coronavirus, Longquan aa coronavirus |
| | | | Small intestine (2) | 28.4% (27/95) | Murine coronavirus, Longquan aa coronavirus, Murine coronavirus \| Longquan aa coronavirus |
| | | | Spleen (1) | 50.0% (1/2) | Murine coronavirus |
| | | | Urine swab (1) | 0% (0/4) | |
| | | Trader | Brain (1) | 4.0% (1/25) | Murine coronavirus |
| | | | Lung (4) | 12.8% (6/47) | Murine coronavirus, Murine coronavirus \| Longquan Aa mouse coronavirus |
| | | | Oral swab (8) | 18.2% (28/154) | Murine coronavirus |
| | | | Small intestine (7) | 12.1% (14/116) | Murine coronavirus, Longquan aa coronavirus |
| *Rattus* | Wildlife farm | | Environmental sample (1) | 100% (1/1) | Bat coronavirus 512/2005 |
| *Hystrix* | Wildlife farm | | Environmental sample (4) | 7.1% (2/28) | Bat coronavirus 512/2005 |
| | | | Feces (23) | 6.0% (18/299) | Bat coronavirus 512/2005, Infectious bronchitis virus (IBV) |
| | | | Urine swab (3) | 0% (0/4) | |

| | | | | | |
|---|---|---|---|---|---|
| *Rhizomys* | Wildlife farm | | Feces (11) | 6.3% (6/96) | Bat coronavirus 512/2005, Infectious bronchitis virus (IBV) |
| *Sciuridae* | Wildlife farm | | Feces (1) | 0% (0/1) | |

[a] Suborder
