## Supplementary material for "Coronavirus testing indicates transmission risk increases along wildlife supply chains for human consumption in Viet Nam, 2013-2014": S2 Table

**S2 Table: Multivariate mixed effect logistic regression showing the association between season and interface with coronavirus positives in field rats in the rodent trade.**

| Variables | Categories | Odds Ratio | 95% CI | P-value |
|---|---|---|---|---|
| Season | Dry | 1.0 | | |
| | Wet | 4.9 | 1.4 - 18.0 | <0.01 |
| Interface | Trade | 1.0 | | |
| | Large market | 2.2 | 1.0 - 4.8 | 0.0326 |
| | Restaurant | 10.0 | 2.7 - 39.5 | <0.01 |

Site was used as grouping variable (i.e. random effect).
