## Supplementary material for "Coronavirus testing indicates transmission risk increases along wildlife supply chains for human consumption in Viet Nam, 2013-2014": S3 Table

**S3 Table: GenBank accession numbers for coronavirus sequences detected in this study and for reference sequences**

| | Sequence Name | Country | GenBank accession number |
|---|---|---|---|
| 1 | Coronavirus_PREDICT_CoV-17/VN13P0020 | Viet Nam | KX285719 |
| 2 | Coronavirus_PREDICT_CoV-17/PB015 | Nepal | KX284941 |
| 3 | Coronavirus_PREDICT_CoV-17/VN13P0024 | Viet Nam | KX285720 |
| 4 | Coronavirus_PREDICT_CoV-17/VN13F0063 | Viet Nam | KX285605 |
| 5 | Coronavirus_PREDICT_CoV-35/VN13F0234 | Viet Nam | KX285678 |
| 6 | Coronavirus_PREDICT_CoV-35/VN13F0226 | Viet Nam | KX285672 |
| 7 | Scotophilus_bat_coronavirus_512_2005/PREDICT-VN13F0113 | Viet Nam | KX285631 |
| 8 | Bat_coronavirus_isolate_BtCoV/B551005/Pte_lyl/CB3-THA/Sep12 | Thailand | MG256467 |
| 9 | Coronavirus_PREDICT_CoV-35/KHP12PTR1-0160B | Cambodia | KX285751 |
| 10 | Coronavirus_PREDICT_CoV-35/GVF-CM-ECO70504 | Cameroon | KX284991 |
| 11 | Coronavirus_PREDICT_CoV-35/CD115889 | DRC | KX285074 |
| 12 | Scotophilus_bat_coronavirus_512_2005/PREDICT-VN13F0276 | Viet Nam | KX285716 |
| 13 | Scotophilus_bat_coronavirus_512_2005/PREDICT-VN13F0161 | Viet Nam | KX285642 |
| 14 | Bat_coronavirus_(BtCoV/A535/2005) | China | DQ648824 |
| 15 | Miniopterus_fuliginosus_coronavirus_isolate_CYCU-M22/TW/2013 | Taiwan | KT381920 |
| 16 | Rhinolophus_monoceros_coronavirus_isolate_CYCU-R14/TW/2013 | Taiwan | KT381916 |
| 17 | Bat_coronavirus_HKU6_PREDICT-EHA-156-12-LS13721 | China | KX285184 |
| 18 | Porcine_epidemic_diarrhea_virus_isolate_KNU-1708 | South Korea | MH052687 |
| 19 | Human_coronavirus_229E | DRC | KX286258 |
| 20 | NL63-related_bat_coronavirus_strain_BtKYNL63-9a | Kenya | NC_032107 |
| 21 | Human_coronavirus_HKU1_isolate_SI17244 | Thailand | MH940245 |
| 22 | Rousettus_bat_coronavirus_HKU9_PREDICT-KHP13-BN1-0003 | Cambodia | KX285758 |
| 23 | Murine_coronavirus_KHP13-BLM2-0001 | Cambodia | KX285756 |
| 24 | Bovine_coronavirus_isolate_BCoV/SLO/5580/2013 | Slovenia | KX059621 |
| 25 | Betacoronavirus_HKU24_strain_HKU24-R05010I | China | KM349744 |
| 26 | Human_coronavirus_OC43_isolate_TNP_12643 | Cote d'Ivoire | MG977452 |
| 27 | Night-heron coronavirus HKU19 | China: Hong Kong | NC_016994 |
| 28 | Bat_coronavirus_isolate_BtCoV/B55080/S.hea/CB/Tha/1/2012 | Thailand | KJ020603 |
| 29 | SARS-related_coronavirus_isolate_F23 | China | KU973688 |
| 30 | Bat coronavirus HKU2 isolate HKU2-1 | Hong Kong | DQ249235 |
| 31 | Bat coronavirus HKU4 isolate HKU4-4 | Hong Kong | DQ249216 |
| 32 | Bat coronavirus HKU8 isolate HKU8-1 | Hong Kong | DQ249228 |
| 33 | Rousettus bat coronavirus HKU10 isolate 183A | China | JQ989270 |
| 34 | Coronavirus PREDICT CoV-35/VN13F0060 | Viet Nam | KX285602 |
| 35 | Scotophilus bat coronavirus 512 2005/PREDICT-VN13F0065_F1R | Viet Nam | KX285606 |
| 36 | Murine_Coronavirus_PREDICT-VN13M0058-T3R | Viet Nam | MT221700 |
| 37 | Murine_Coronavirus_PREDICT-VN13M0002-O1R | Viet Nam | MT221698 |
| 38 | Longquan aa coronavirus PREDICT-VN13M0299-T1R | Viet Nam | MT221699 |
| 39 | Longquan aa coronavirus_PREDICT-VN13M0005-F1R | Viet Nam | MT221701 |

| | | | |
|---|---|---|---|
| 40 | Scotophilus_bat_coronavirus_512_2005/PREDICT-VN14F0283_F1R | Viet Nam | MT221704 |
| 41 | Scotophilus_bat_coronavirus_512_2005/PREDICT-VN14F0014_E1R | Viet Nam | MT221703 |
| 42 | Scotophilus_bat_coronavirus_512_2005/PREDICT-VN13F0333_E1R | Viet Nam | MT221702 |
| 43 | Coronaviridae_sp._isolate_75-990L01-R5-CoV-BAT-VN | Viet Nam | KX092205 |
| 44 | Coronaviridae_sp._isolate_75-55-L01-R4-CoV-BAT-VN | Viet Nam | KX092189 |
| 45 | Coronaviridae_sp._isolate_75-98-L07-R4-CoVBAT-VN | Viet Nam | KX092190 |
| 46 | Coronaviridae_sp._isolate_75-98-L07-R2-CoV-BAT-VN | Viet Nam | KX092177 |
| 47 | Coronaviridae_sp._isolate_7562L01R6RATCoV | Viet Nam | KX092217 |
| 48 | Coronaviridae_sp._isolate_7565L07R5RATCoV | Viet Nam | KX092223 |
