## Supplementary material for "Coronavirus testing indicates transmission risk increases along wildlife supply chains for human consumption in Viet Nam, 2013-2014": Author Contributions

Funding acquisition: Jonna A. K. Mazet, Damien O. Joly

Investigation: Nguyen Thi Thanh Nga, Nguyen Van Long (WCS), Amanda E. Fine, Nguyen Tung, Van Dang Ky, Phan Quang Minh, Nguyen Thi Diep

Lab testing, interpretation and analysis: Le Tin Vinh Quang, Nguyen Thi Hoa, Tracey Goldstein, Alexandre Tremeau-Bravard, Victoria Ontiveros

Project administration: Nguyen Van Long (DAH), Nguyen Thi Thanh Nga, Nguyen Van Long (WCS), Amanda E. Fine, Sarah H. Olson, Jonna A. K. Mazet, Christine Kreuder Johnson, Tracey Goldstein, Damien O. Joly, Martin Gilbert, Leanne Wicker

Supervision: Scott I. Roberton, Amanda E. Fine, Damien O. Joly, Chris Walzer, Hoang Bich Thuy, Nguyen Van Long (DAH), Nguyen Thanh Phuong, Bach Duc Luu, Vo Van Hung, Nguyen Thi Lan

Writing – original draft: Nguyen Quynh Huong, Mathieu Pruvot, Amanda E. Fine, Nguyen Thi Thanh Nga, Alice Lattine, Sarah H. Olson

Writing – review & editing: All
